# A method to minimize condenser lens-induced hysteresis effects in a JEOL JEM-3200FSC microscope to enable stable cryoEM low-dose operations

**DOI:** 10.1101/153395

**Authors:** M. Jason de la Cruz, Michael Martynowycz, Johan Hattne, Dan Shi, Tamir Gonen

## Abstract

Low dose imaging procedures are key for a successful cryoEM experiment (whether by electron cryotomography, single particle analysis, electron crystallography, or MicroED). We present a method to minimize magnetic hysteresis of the condenser lens system in the JEOL JEM-3200FSC transmission electron microscope (TEM) in order to maintain a stable optical axis for the beam path of low-dose imaging. The simple procedure involves independent voltage ramping of the CL1 and CL2 lenses immediately before switching to the focusing and exposure beam settings for data collection.

**Highlights:**

- Enables consistent horizontal beam position during low-dose mode switching
- Facilitates EM automated data collection for the JEOL JEM-3200FSC
- Improves beam stability in both TEM imaging and diffraction modes

## 1. Introduction

CryoEM low-dose experiments place much stress on a TEM’s electromagnetic lenses due to the frequent changes necessary to alter the magnetic fields for changing the electron beam’s trajectory down the optical axis of the microscope. For each data collection session, the same beam must be used for multimodal imaging, each with different beam settings: low-magnification searching, high-magnification focusing, tracking methods, and final exposure. In automated data collection schemes, many cycles of these modes iterate in order to collect many different areas on a sample grid. Therefore, a quickly-stabilized beam is desirable.

Magnetic hysteresis affecting beam position in a transmission electron microscope (TEM) is well known and has been documented using the JEOL JEM-3200FSC [1]. Hysteresis is an intrinsic property of many magnetic materials and is present throughout the electron optics of a TEM, for example, in the different lens systems, deflection coils, pole piece, and energy filter (if so equipped). We have found that the main contributor to inconsistencies in beam movement and position upon changing between different lens programs for cryoEM data collection purposes on the JEOL JEM-3200FSC is magnetic hysteresis caused by the condenser lens system.

Magnetic hysteresis refers to the persistent, atomic alignment of a ferromagnetic material’s magnetic dipole moments in the presence of an external magnetic field. Hysteresis is evident by the retention of magnetization after the external field has been removed. The presence of magnetic domains, or sub-units, within the soft magnetic material of the lenses may impart their own directionality upon magnetism in a particular direction by electrical current. Upon initial excitation to change the directionality of the magnetic field, not all magnetic domains will initially orient in the desired direction. Therefore, in order to orient the directionality of all the magnetic domains in the magnetic material in the same direction, more energy is necessary to drive the soft magnet to saturation [2].

Current methods to “correct” these lens effects are not suitable for the rapid lens program switching required for automated data collection or cryoEM applications, due to the time and manual effort needed to adequately stabilize the beam. Other microscopes, including models from electron microscope manufacturers JEOL and FEI, have proprietary normalization schemes and lens designs optimized to minimize beam movement due to magnetic hysteresis. We describe a robust method to normalize the condenser lens system for the JEOL JEM-3200FSC microscope, enabling efficient and automated low-dose cryoEM data collection.

## 2. Magnetic hysteresis in the condenser lens

Several electromagnetic lenses in a TEM can contribute to beam instability during lens mode switching during automated data collection procedures. However, magnetic hysteresis due to the condenser lens system has the greatest influence because, as the first lens downstream from the electron gun, it controls primary illumination of the sample.

The condenser lens in a TEM is made up of ferromagnetic material specifically arranged within an annular structure that makes up part of the microscope column [***Fig. 1***]. The type of soft magnetic material used to construct the lens, as well as its design in the microscope, can affect the stability of the magnetic fields, and hence the beam on the specimen. Modern TEMs typically have two such lenses, and the user can generally control both. Adjusting the voltage to each of these lenses alters the focal length of the beam crossover that follows each lens [3]. The first condenser lens, CL1, lies close to the electron source with only the gun deflector coils separating them. The CL1 lens condenses, or demagnifies, a focused image of the source onto a plane somewhere before the second condenser lens “CL2” (actually termed “CL3” in the JEOL JEM-3200FSC PC interface) and condenser aperture. Excitation of the CL1 lens causes beam crossover above the aperture, thus spreading the beam at the level of CL2 and causing a decrease in beam current and intensity. This initial step-down of the beam current is known as setting the “spot size” or beam diameter.

**Figure 1.**
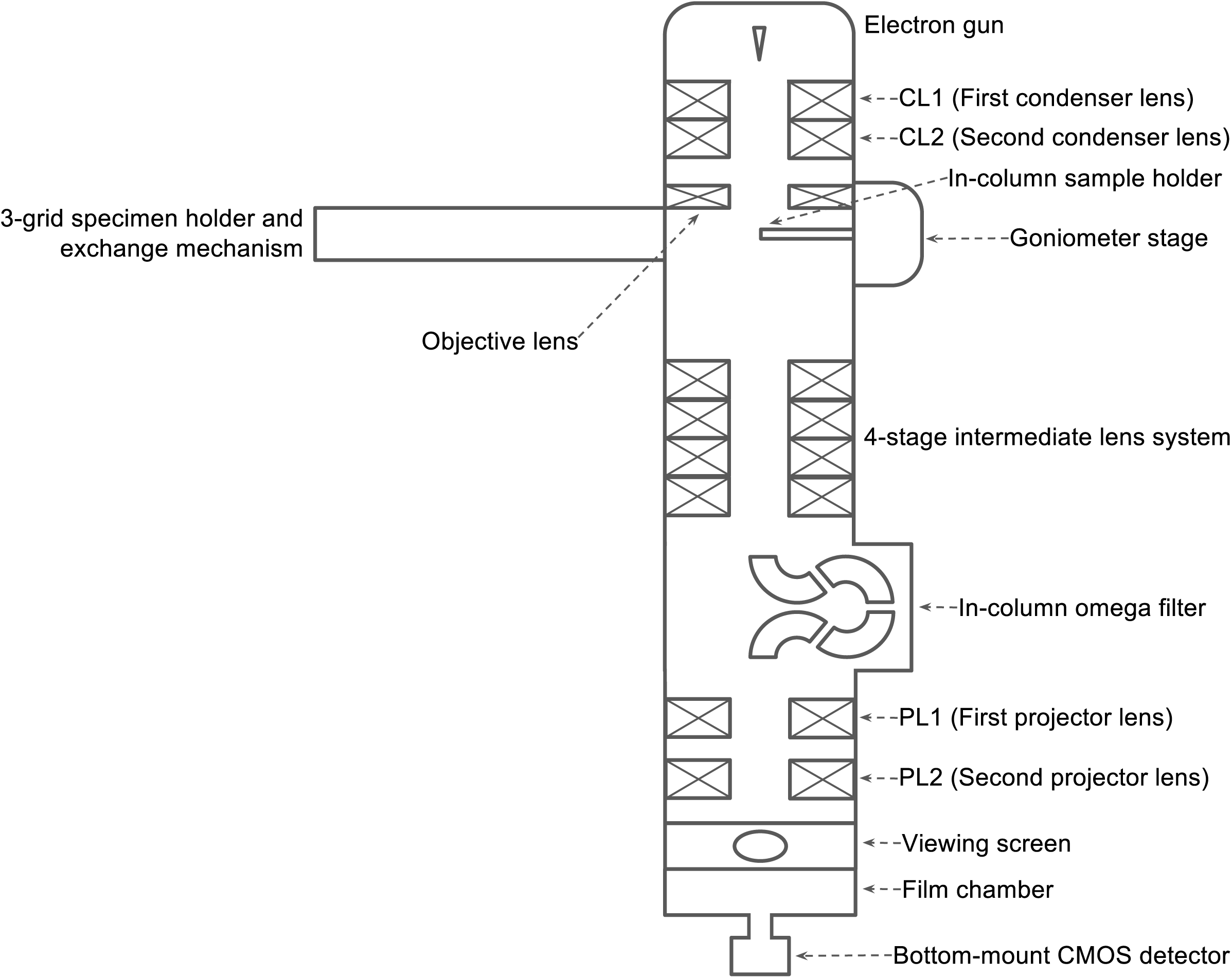
Schematic diagram of lenses in the JEOL JEM-3200FSC TEM at the Janelia Research Campus.

Fine-tuning of the beam current at the specimen plane involves the adjustment of the CL2 voltage. On the microscope, the “BRIGHTNESS” knob alters the CL2 value, which adjusts the focal length of beam crossover to above or below the specimen. For cryoEM imaging and diffraction experiments, beam convergence should be at a minimum (*i.e.*, the beam should be focused at the specimen plane) to ensure imaging at or close to parallel illumination [4].

## 3. Low-dose data collection

Low-dose data collection is essential for cryoEM methods in order to reduce the energy deposited to the sample [5]. For data collection schemes requiring the use of the objective lens (*i.e.*, TEM imaging), three lens programs are defined with their own spot size, intensity, magnification, focus, and other lens and deflector settings: Search mode, Focus mode, and Exposure mode. Users tend to collect data inside patterned TEM grid foil holes of certain size. In Search mode, the beam is generally set to lowest magnification in the microscope’s main upper magnification range, and defocused enough to see the centered targeted hole and surrounding carbon areas for focusing. Focus mode tilts the beam to an off-center area and is set at the same magnification as exposure mode or higher. The focus mode can also be set to image the focusing area in defocused-diffraction mode which applies less total electron dose per unit area than its equivalent area and illumination conditions in imaging mode [1]. The beam position in exposure mode should be at the optical axis for data collection.

Low-dose techniques are also necessary for MicroED [6,7] data collection in order to minimize pre-irradiation of the crystal. Two of the lens programs are necessary for MicroED: Focus mode and Exposure mode. Focus mode sets the beam in highly-defocused diffraction mode for crystal positioning and set-up for data collection. In Exposure mode, the beam is set in diffraction mode with minimum convergence at the specimen plane. An optional Search mode can be used for searching grid squares at low magnification, but it is not strictly necessary.

## 4. Experimental Procedures

Empty R2/2 Quantifoil [8] TEM grids served as a sample substrate for these experiments and were examined using a JEOL JEM-3200FSC field emission TEM operated at 300 kV and equipped with an in-column omega energy filter. Images were recorded using a bottom-mounted TVIPS TemCam-F416 CMOS detector controlled by SerialEM 3.6.0. SerialEM is a third-party data acquisition program for electron microscopes [9]. To demonstrate beam stability along the optical axis with this procedure, we simulated lens program switching as it applies to single particle/tomography data collection and MicroED.

The JEOL JEM-3200FSC PC interface includes a module, the FLC Panel (for Free Lens Control), for manual control of the voltage for each lens in the system [***Fig. 2***]. The units are displayed with 16-bit precision either in hexadecimal notation or unsigned 5-digit decimal form to represent a percentage of the maximum current. The lens controls have an “on” and “off” state, because these controls override the standard system settings. Normal operation of the microscope requires that all FLC Panel controls be in the “off” state. Changes enabled here will turn the selected lens “on”. In this panel, the second condenser lens is called “CL3”.

**Figure 2.**
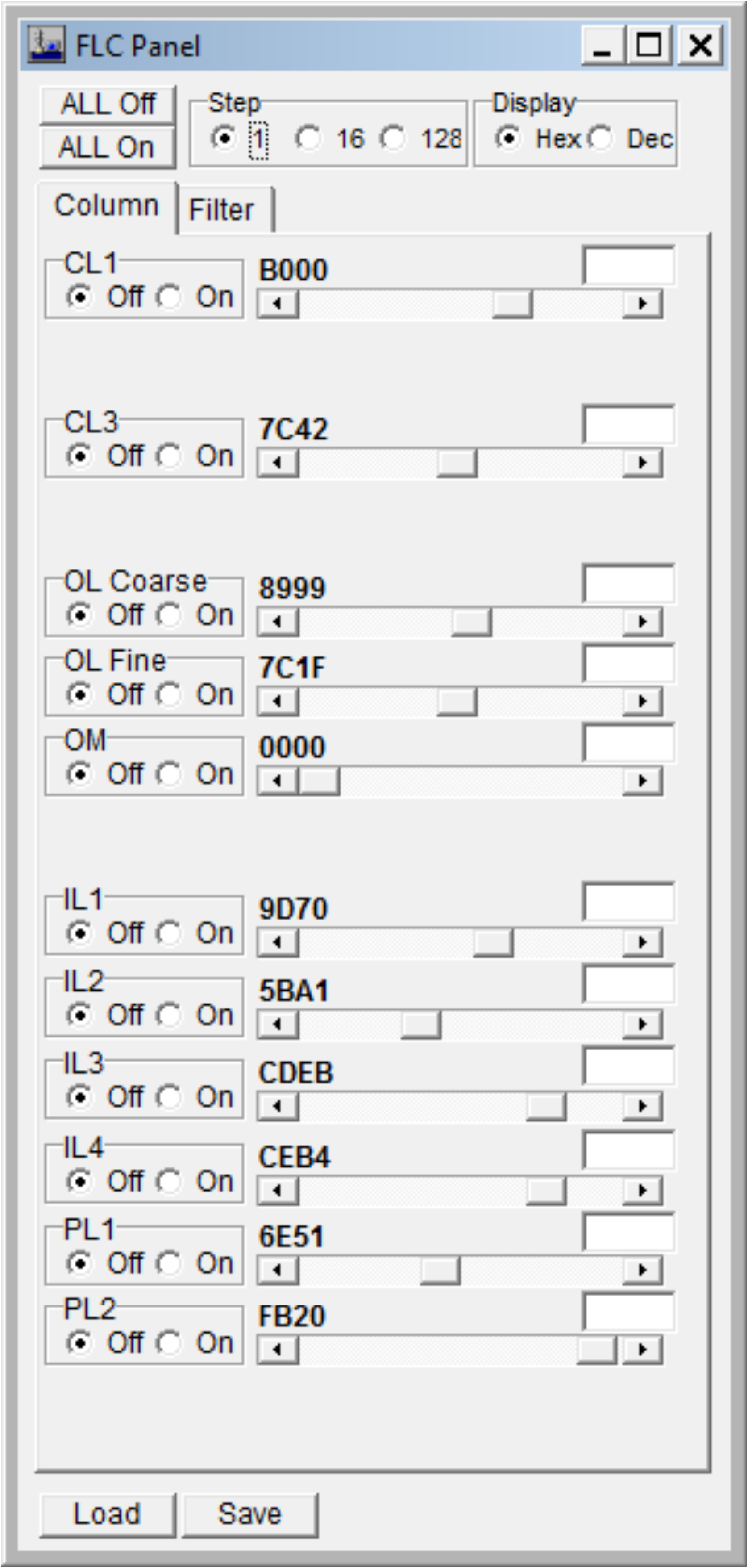
JEOL PC interface dialog window: Free Lens Control panel.

The normalization procedure involves only the first (CL1) and second (CL2) condenser lenses, with the CL1 lens done first. The current is ramped to its maximum setting, and then returned to its previous setting. This is performed twice, in quick succession, for each lens. The normalization procedure is invoked immediately before switching the lens program to Focus mode or Exposure mode. In the FLC Panel, the user simply slides the switch for CL1 completely to the right, releases the button so the lens value changes to maximum applied current, then presses the “off” radio button for CL1 to the left, returning the lens to the previous value. This is done again for CL1. Then the procedure is repeated for CL2. Once finished, the beam is ready for subsequent use.

In our trials, before starting an imaging test, the microscope was set to diffraction settings, and vice-versa. This was done to “reset” the condenser lens to different conditions to avoid magnetic memory. Also, for the imaging experiments, the beam was condensed so that the edges can be seen, as we are tracking beam movement as a test for stability.

## 5. Discussion

A simple method to compensate for magnetic hysteresis in the condenser lens system of the JEOL JEM-3200FSC TEM has been described, by increasing voltage of the condenser lenses to the maximum value (ramping) twice, each in succession, from the gun area towards the specimen. The iterative ramping ensures that the magnetic domains within the ferromagnetic core of the lenses are pointing in the same direction after lens mode switching; this in turn leads to a more stable beam recall to previous lens and deflector settings and beam position stability. The sequence of the normalization procedure is important because the CL1 and CL2 lenses emit strong magnetic fields, even external to the microscope. Due to the close proximity of the lenses to each other, a strong change to the magnetic field in one lens will affect the magnetic field of the other by magnetic induction. Therefore, the sequence in normalizing the condenser lenses follows the path of the electron beam: CL1, then CL2.

Other lenses are present in the microscope, for example the objective and projector lens systems. However, beam stability in terms of recall and short-term positioning are not affected in those lenses. In addition, our experiments showed that this normalization procedure benefits imaging mode more than diffraction mode, likely due to the additional lenses involved in imaging mode. In all images involving a condensed beam, the initial setting was set to the center of the image. Also, for both imaging and diffraction experiments, a single cycle between Focus and Exposure modes were done before the final Exposure image was taken. For imaging mode, normalization was not required for Search mode because the beam in this mode was set to low magnification, therefore the observed beam shift was negligible [***Fig. 3A, D***]. Normalization was found to be beneficial prior to switching between Focus and Exposure modes, keeping the beam at the center of the screen [***Fig. 3B-C, E-F***]. In diffraction mode on our microscope, normalization was not required for the defocused-diffraction Focus mode [***Fig. 4A, C***] but required for Exposure mode [***Fig. 4B, D***].

**Figure 3.**
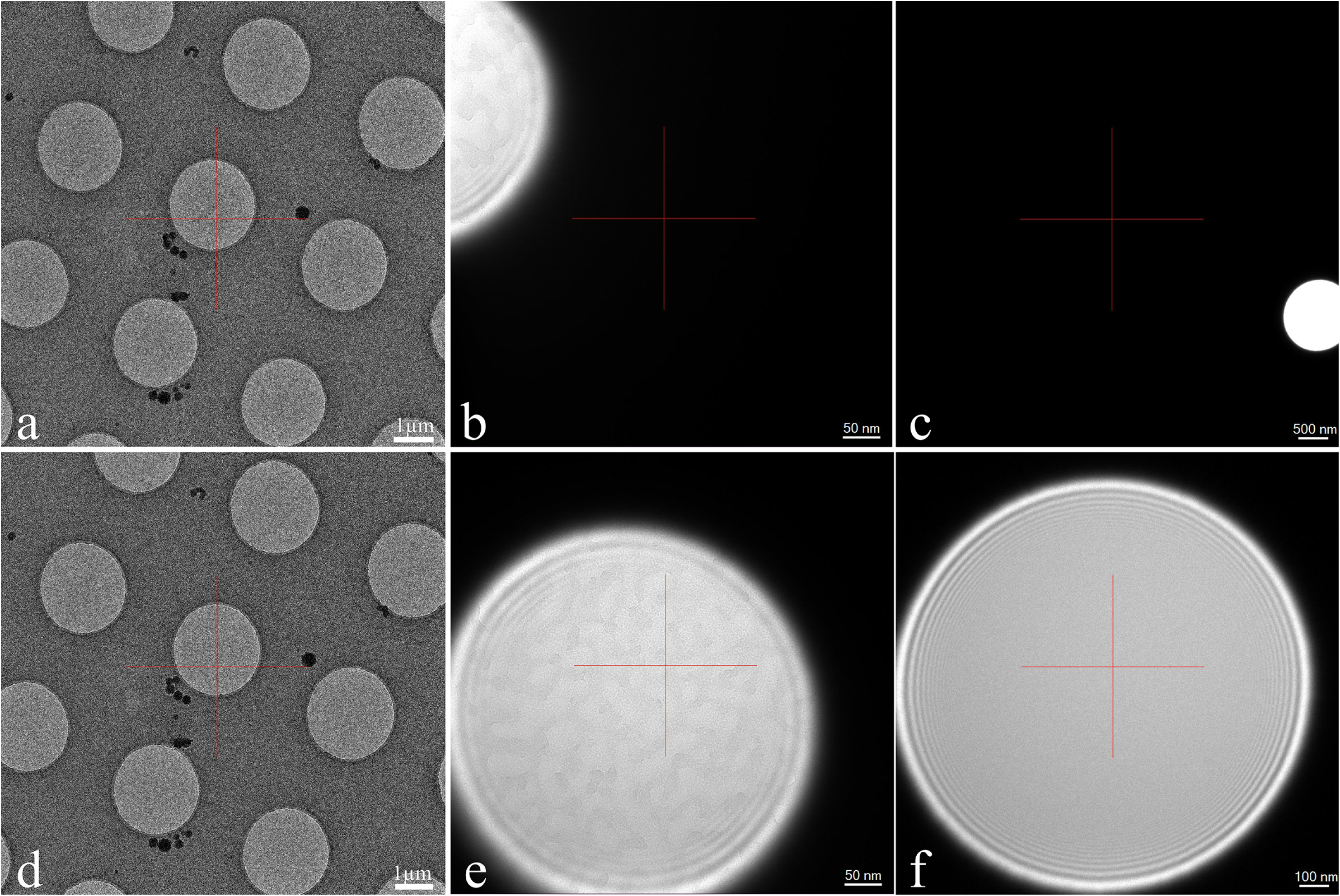
Beam stability in imaging mode: lens program for single particle data collection. (A) Search mode without normalization: magnification 2 500 X, spot 6, CL3 excitation 55.24%. (B) Focus mode without normalization: magnification 50 000 X, spot 5, CL3 excitation 50.31%. (C) Exposure mode without normalization: magnification 4 000 X, spot 8, CL3 excitation 48.58%. Here, the beam had shifted off the detector’s field of view at the original magnification of 20 000 X; this image shows rotation-free demagnification of the area at 4 000 X. (D) Search mode without normalization: magnification 2 500 X, spot 6, CL3 excitation 55.24%. (E) Focus mode with normalization: magnification 50 000 X, spot 5, CL3 excitation 50.31%. (F) Exposure mode with normalization: magnification 20 000 X, spot 8, CL3 excitation 48.56%.

**Figure 4.**
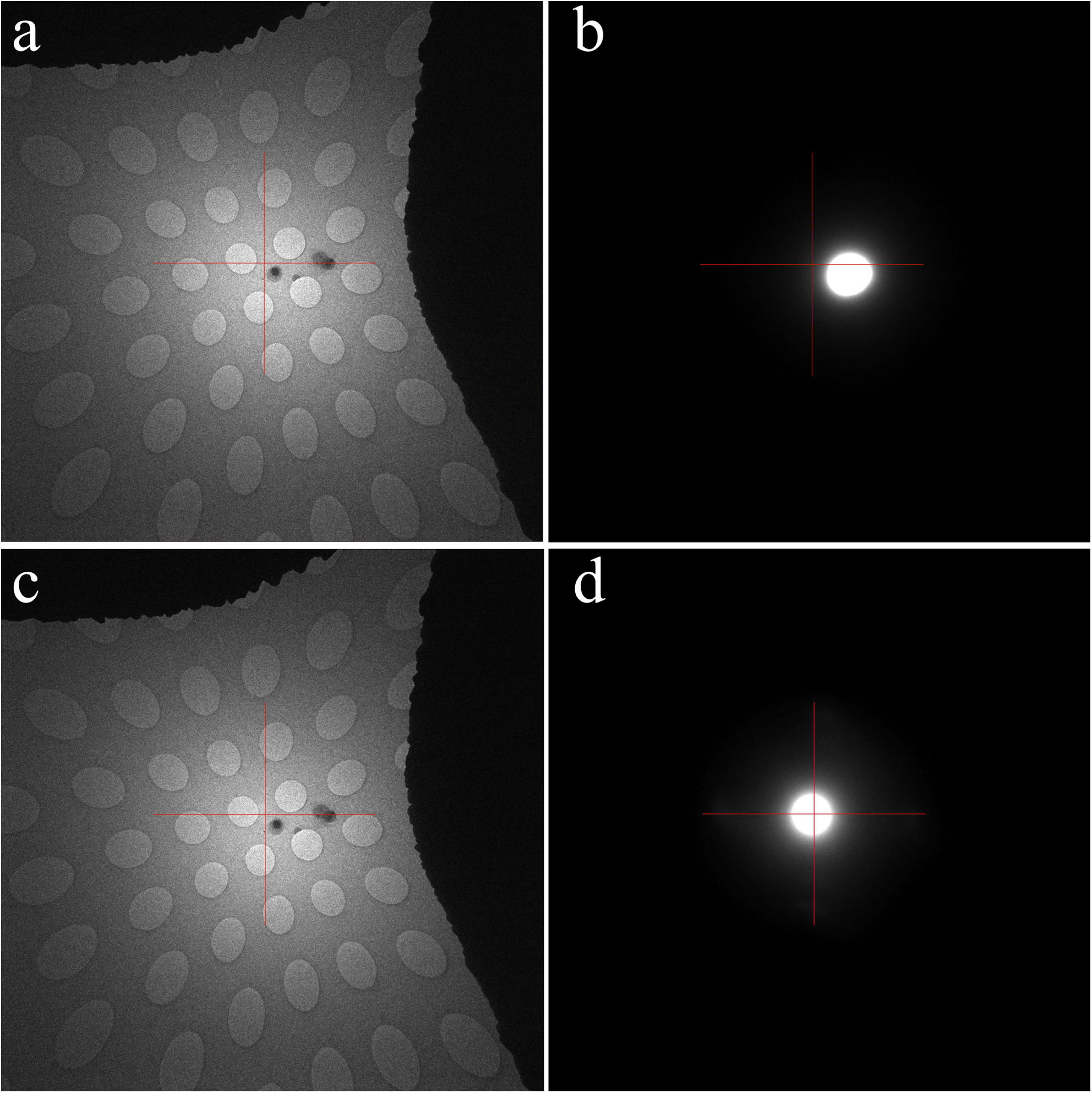
Diffraction beam stability. (A) Focus mode without normalization: camera length 30 m, spot 8, CL3 excitation 59.6%. (B) Photo/exposure mode without normalization: camera length 120 m, spot 8, CL3 excitation 48.79%. (C) Focus mode with normalization: camera length 30 m, spot 8, CL3 excitation 59.6%. (D) Photo/Exposure mode with normalization: camera length 120 m, spot 8, CL3 excitation 48.79%.

Other methods of compensating for magnetic hysteresis include: resetting the condenser lens and activating the beam tilt wobbler [10] and lens relaxation by current oscillation [4, 11], among others. However, these are lengthy procedures that are intractable to high-throughput serial data collection in cryoEM. Another approach to overcome magnetic hysteresis increases the beam diameter beyond the edges of the detector area, exposing nearby areas on the grid to extra dose. For cryoEM data collection, the use of a large beam diameter, with additional room allowed for beam drift, is not preferable because it would limit the number of areas one can collect on a single grid. Another method utilizes stage memory to save the goniometer coordinates to memory, fixes the beam parameters in another area of the grid, and then goes back to the saved position for exposure. Stage recall, however, is generally not precise enough for high magnification imaging and usually requires additional adjustment prior to each exposure.

The major advantage of condenser lens normalization using this method is stability of specimen illumination throughout lens mode switching in automated and cryoEM data collection. By promoting the stability of the condenser lens system, one can ensure that previously-set imaging conditions are recalled, especially to ensure parallel illumination of the sample. The normalization procedure is quickly done manually, and can be scripted for automatic application.

## Acknowledgements

The authors are grateful for the expertise and technical support of Reza Ghadimi (TVIPS GmbH) and Sohei Motoki (JEOL Ltd.). We would like to thank Oliver Hofnagel (Max Planck Institute of Molecular Physiology) for testing this method on the JEM3200FSC at the MPI in Dortmund. We also thank Zhiheng Yu (HHMI/Janelia Research Campus) for critical reading of the manuscript and Ruben Diaz-Avalos (HHMI/Janelia Research Campus) for insightful discussions. The Gonen laboratory is supported by funds from the Howard Hughes Medical Institute.

